# A rapid, point of care red blood cell agglutination assay for detecting antibodies against SARS-CoV-2

**DOI:** 10.1101/2020.05.13.094490

**Authors:** Robert L. Kruse, Yuting Huang, Heather Smetana, Eric A. Gehrie, Tim K. Amukele, Aaron A.R. Tobian, Heba H. Mostafa, Zack Z. Wang

## Abstract

The COVID-19 pandemic has brought the world to a halt, with cases observed around the globe causing significant mortality. There is an urgent need for serological tests to detect antibodies against SARS-CoV-2, which could be used to assess the prevalence of infection, as well as ascertain individuals who may be protected from future infection. Current serological tests developed for SARS-CoV-2 rely on traditional technologies such as enzyme-linked immunosorbent assays (ELISA) and lateral flow assays, which may lack scalability to meet the demand of hundreds of millions of antibody tests in the coming year. Herein, we present an alternative method of antibody testing that just depends on one protein reagent being added to patient serum/plasma or whole blood and a short five-minute assay time. A novel fusion protein was designed that binds red blood cells (RBC) via a single-chain variable fragment (scFv) against the H antigen and displays the receptor-binding domain (RBD) of SARS-CoV-2 spike protein on the surface of RBCs. Upon mixing of the fusion protein, RBD-scFv with recovered COVID-19 patient serum and RBCs, we observed agglutination of RBCs, indicating the patient developed antibodies against SARS-CoV-2 RBD. Given that the test uses methods routinely used in hospital clinical labs across the world, we anticipate the test can be rapidly deployed with only the protein reagent required at projected manufacturing cost at U.S. cents per test. We anticipate our agglutination assay may find extensive use in low-resource settings for detecting SARS-CoV-2 antibodies.

## Introduction

The SARS-CoV-2 coronavirus causing COVID-19 disease represents growing pandemic, leading to acute respiratory distress syndrome in a small portion of patients and ultimately significant mortality.^1^ SARS-CoV-2 imposes both diagnostic and therapeutic challenges. On the diagnostic front, reverse transcriptase-polymerase chain reaction (RT-PCR) is the gold standard for detecting the virus, but PCR represents significant costs in low-income countries and is not always widely available.^2^ Importantly, RT-PCR cannot detect evidence of past infection, which will be crucial for epidemiological efforts to assess how many people have been infected. While viral load determined by RT-PCR appears to have prognostic value,^3^ measuring the immune response against SARS-CoV-2 may also be beneficial into assessing the outcomes of patients and understanding global prevalence.^4^

Serologic testing for antibodies against SARS-CoV-2 could detect both recent and past infection, which is crucial for surveillance and epidemiological studies.^5^ However, current enzyme-linked immunosorbent assay (ELISA) tests for COVID-19 require a number of steps, washes, and reagents, involving hours of manual time and/or automated machines.^6^ Lateral flow immunoassays have been developed for SARS-CoV-2, but still require the manufacturing of strips, plastic holders, and multiple different antibody types and conjugates.^7^ A rapid lateral flow immunoassay is also limited as one-time use. There is an urgent need for a low complexity assay that could be performed as a point of care test in low-income countries, without the need for machines.

As an alternative method to detect antibodies, blood banks across the world routinely detect antibodies against blood group antigens as part of the type and screen assay, performed before blood transfusions are given. The readout of the assay is hemagglutination, or the aggregation of red blood cells (RBCs), which can be captured by a camera or easily observed with the naked eye. Furthermore, hemagglutination testing, whether by hand or by an automated machine, can be used to titer antibodies, measuring their levels in the serum.^8^ This particular flexibility to range from point of care, single patient testing to wide scalable on existing automated platforms in clinical labs is unique among the different serologic options.

Hemagglutination has been leveraged in the past to detect antibodies against pathogens. The first iteration consisted of cross-linking an antibody against RBC antigens with a peptide antigen from human immunodeficiency virus (HIV).^9,10^ When incubated with whole blood from HIV patients, RBC agglutination could be observed, indicating antibodies specific to that antigen were detected.^9^ A comparison of 1800 patient blood specimens found similar sensitivity and specificity among commercial ELISA kits and 2-minute autologous RBC agglutination testing.^11^ Later studies improved on the technology by building fusion proteins of antibody fragments with antigens from HIV.^12,13^ Targeting multiple different RBC antigens at the same time improved performance characteristics of the assay.^14^ Antibodies against West Nile virus have also been detected by autologous RBC agglutination assay.^15^ Outside of infectious disease, elevated D-dimer levels could also be detected with a similar red agglutination assay, SimpliRED, for point of care testing for patients with suspected deep vein thrombosis.^16,17^

Herein, we describe an RBC agglutination assay to detect antibodies against the receptor-binding domain (RBD) of SARS-CoV-2 spike protein in COVID-19 patients, which is the frequent target of neutralizing antibodies against coronaviruses.^18^ The assay may find use in low-resource settings as a rapid method of testing for current or past SARS-CoV-2 infection.

## Methods

### Gene construction

A fusion protein of SARS-CoV-2 antigen fused to a single-chain variable fragment (scFv) targeting an RBC antigen was constructed (**Figure 1**). Briefly, the RBD of the SARS-CoV-2, consisting of amino acids 330 – 524 of the spike protein^19^ was connected via a linker to an scFv derived from the antibody 2E8 targeting the H antigen on RBCs^20^ to form RBD-scFv. RBD-scFv also contained an IgG heavy chain secretion signal to allow export from mammalian cells, and a hexa-histidine tag located at the end of the protein to allow for convenient purification. The RBD-scFv was synthesized (Twist Bioscience) and cloned into pIRII-IRES-GFP^21^ to form pIRII-RBD-scFv-IRES-GFP.

**Figure 1.**
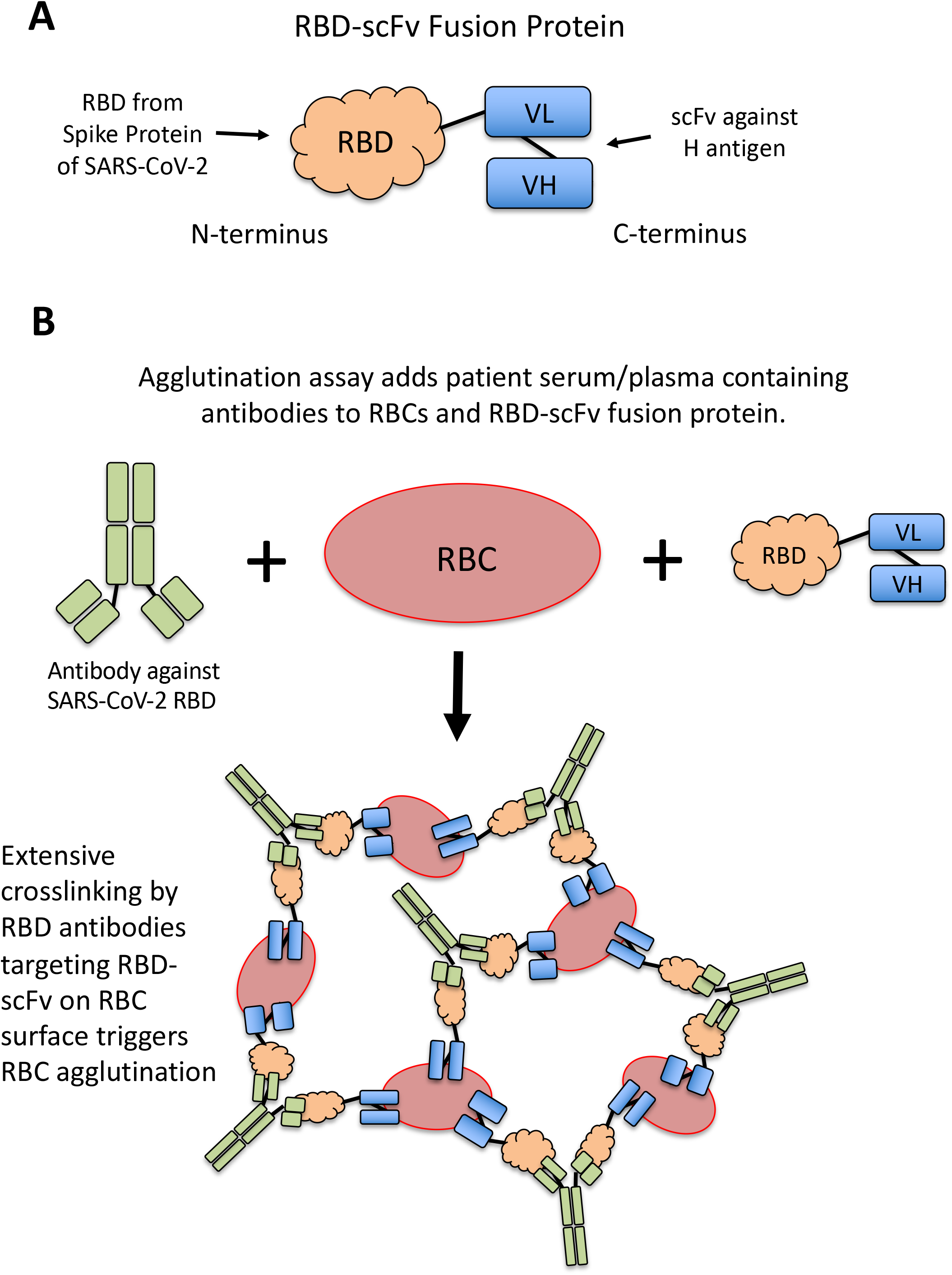
Construction of fusion protein and mechanism of agglutination. (A) A fusion protein consisting of the receptor-binding domain (RBD) of the SARS-CoV-2 spike protein at the N-terminus connected via a linker to a single-chain variable fragment (scFv, consisting of VH and VL domains connected with a flexible linker) at the C-terminus targeting the H antigen on the surface of red blood cells (RBCs). (B) Patient serum/plasma containing antibodies targeting the RBD are mixed with RBCs and RBD-scFv fusion protein and allowed to cross-link multiple RBCs in mass to eventually lead to visible agglutination seen with the naked eye. VH = variable heavy; VL = variable light

### Protein Production and Purification

RBD-scFv protein was produced in HEK 293T cells in order to preserve proper folding and glycosylation patterns of the viral domain. pIRII-RBD-scFv-IRES-GFP and pCMV-hyperPB (hyperactive piggyBac transposase) were co-transfected using Lipofectamine 3000 into HEK 293T cells. The piggyBac transposon system afforded stable integration and protein production. The supernatant containing RBD-scFv was collected at 72 hours. RBD-scFv was purified from the supernatant using Capturem™ His-Tagged Purification Miniprep Kit (Takara Bio). One prep of 800 μL supernatant through one column of the kit yielded 102 μg/mL of RBD-scFv measured by NanoDrop™ 2000/2000c Spectrophotometers (ThermoFisher), which was used in subsequent experiments. The correct size of the purified RBD-scFv protein (52.4 kDa → ∼57 kDa with glycosylation) was confirmed by protein gel electrophoresis using Mini-PROTEAN TGX Precast Gels 4-20% (Bio-RAD).

### Red blood cell agglutination testing

A deidentified, discarded serum sample from a recovered COVID19 patient was evaluated. The patient specimen was collected greater than 28 days post symptoms of COVID-19 with negative PCR testing upon discharge; the patient was previously positive by nasopharyngeal swab PCR testing.

For the RBC agglutination assay, a round bottom 96-well plate was used. O-type Rh-positive red blood cells suspended in 2-4% solution (Immucor) were obtained from the Johns Hopkins Blood Bank. 20 μL of red blood cell solution, 10 μL of undiluted patient serum and 10 μL of RBD-scFv solution were pipetted into each well, following a similar protocol from a previous study.^22^ The solution was thoroughly mixed with the pipette and allowed to incubate for five minutes at room temperature before agglutination was visualized by the naked eye. For testing patient serum, a dilution series of RBD-scFv was performed to test for optimal levels of protein to induce agglutination in presence of patient anti-RBD antibodies. A series of six 1:1 dilutions were performed from the 102 μg/mL RBD-scFv stock. A seventh well of containing patient serum, 10 μL of phosphate-buffered saline (PBS) and RBC solution alone was used as a negative control to rule out potential patient alloantibody induced agglutination, or alternatively, cold reactive IgM autoantibodies.

## Results

### Construction of fusion protein design

The RBD-scFv fusion protein was designed as depicted in **Figure 1A** to decorate SARS-CoV-2 antigens on the surface of RBCs. These decorated RBCs would then yield hemagglutination in the presence of SARS-CoV-2 antibodies (**Figure 1B**). For this proof of concept, the RBD of the SARS-CoV-2 spike protein, corresponding to amino acids 330 – 524 of the spike protein,^19^ was chosen for its small size and stable folding, as well as the fact that the RBD is the target of the majority of neutralizing antibodies against coronaviruses.^23^ Any positive test for antibodies binding to RBD would be highly suggestive of the presence of neutralizing antibodies that would be protective of reinfection.^24^

The scFv connected to the RBD targets the H antigen, which a carbohydrate antigen located within the ABO polysaccharides.^25^ The H antigen is ubiquitous in RBCs in the human population, except among Bombay individuals, who are exceptionally rare.^26^ The H antigen can be confirmed to be expressed using the Ulex europaeus lectin.^27^ A previous study indicated that the scFv 2E8 could successfully bind to RBCs and be used to display HIV gp41 in an assay to detect HIV antibodies in a similar RBC agglutination assay.^22^ The RBD-scFv fusion protein was collected from cell culture supernatant after the transfection of expression plasmids in 293T cells. After purification with a nickel column via His-tag affinity, the RBD-scFv protein was run on a protein gel to confirm the proper size (**Figure 2**).

**Figure 2.**
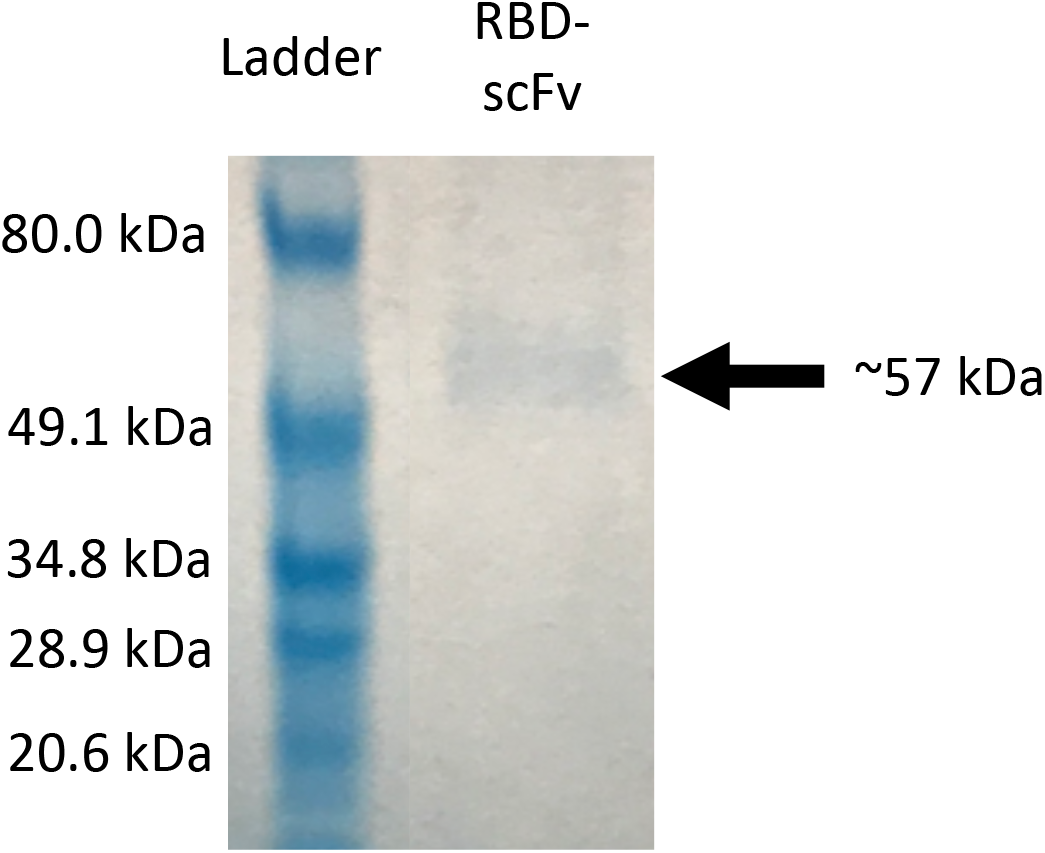
RBD-scFv fusion protein was successfully isolated. RBD-scFv protein was expressed in 293T cells, secreted into the supernatant, and purified by Nickel-His tag affinity columns. Protein gel electrophoresis was performed to confirm the expected band size (∼57 kDa) of the purified RBC-scFv protein.

### Red blood cell agglutination assay to detect SARS-CoV-2 antibodies

The RBD-scFv fusion protein was mixed with RBCs in the presence of COVID19 patient serum in order to detect for agglutination. The assay was carried out in a small total volume, 40 μL, in a 96-well plate. The RBC solution (2-4% red blood cells) is at a dilution commonly used in manual tube testing for ABO typing in blood banks.

Given that the amount of RBD-scFv fusion protein necessary to cross-link antibodies and RBCs and trigger agglutination as unknown, a dilution series was performed, starting with the maximal concentration of the RBD-scFv stock solution through 5 successive 1:1 dilutions. A negative control well contained RBCs and COVID-19 patient serum alone. After five minutes of incubation, agglutination was observed in the three highest concentrations of RBD-scFv protein, with no agglutination observed in the more dilute RBD-scFv concentrations (**Figure 3**). Decreasing levels of agglutination were observed in the three highest levels of RBD-scFv. Incubation of RBD-scFv and RBCs without patient serum did not yield any agglutination (**Figure 3**). This indicates the presence of antibodies that bind to RBD in the patient’s serum, with agglutination activity reflective of RBD-scFv concentration.

**Figure 3.**
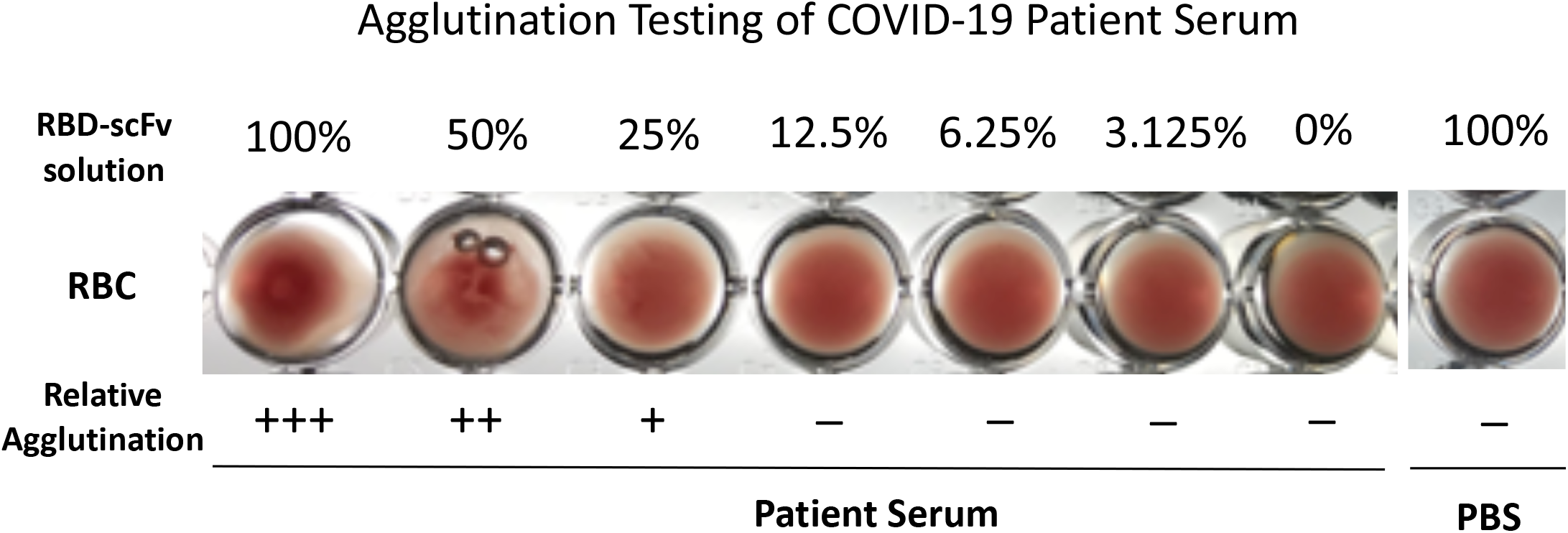
RBD-scFv mediates agglutination the of red blood cells in the presence of COVID-19 patient serum. RBD-scFv protein stock solution (102 μg/mL) was serially diluted in the presence of fixed concentrations of red blood cells (O-negative, 2-4% solution) and undiluted patient serum. Agglutination was seen in the highest three concentrations after five minutes of incubation, most intense in the highest concentration of RBD-scFv. A negative control contained phosphate-buffered solution (PBS) in the place of patient serum.

## Discussion

In this study, we demonstrated a proof of concept for a rapid, point of care RBC agglutination test for SARS-CoV-2 antibodies. Our experiments show that COVID19 patient serum could efficiently agglutinate RBCs in the presence of RBD-scFv within five minutes of incubation. Our ability to detect RBD antibodies in COVID-19 patient serum matches a previous report, which detected RBD antibodies using an ELISA based assay.^28^ Importantly, antibody binding to the SARS-CoV-2 RBD has not shown cross-reactivity with other coronaviruses,^29^ emphasizing its clinical utility. The decreasing agglutination observed in our dilution series matched previous reports, where agglutination scores were related to the amount of fusion protein in the reaction.^30^

The entire hemagglutination assay should be feasible to operate on a drop of whole blood obtained from a patient finger-stick, given only 30 μL of RBCs and serum were used in this experiment. For whole blood assays, RBD-scFv fusion protein alone solution could mixed with whole blood in order to facilitate the assay. Given the costs and complexities of blood specimen processing, this represents a significant advantage. The estimated protein reagent cost for the research assay using small scale production and purification was 25 cents (U.S.) per test, which should be able to be reduced under 1 cent per test with larger-scale protein production and purification. While the assay was carried out in 96 well plates similar to previous studies,^22^ the format is readily transferrable to slide agglutination test, where the components are mixed on a re-usable glass slide by stirring followed by short incubation.^14^ This format is still used for ABO testing in low-resource settings, which suggest ready ability to be deployed. While the point of care assay potential is emphasized in this study for simplicity, the fusion protein reagents could potentially be added into other hemagglutination assays used in clinical labs today, such as tube testing and gel testing, as well as on automated solid-phase machines. The potential repurposing of automated hemagglutination assay machines, in particular, could allow for high-throughput testing of hundreds of thousands of tests today using existing clinical infrastructure.

A key limitation of this assay is that it does not distinguish between IgG, IgA, or IgM against SARS-CoV-2, which may be desired in certain clinical scenarios. IgG subclasses can similarly not be distinguished. Also, the assay is highly dependent on the viral antigen deployed, such that clinicians should be aware if RBD antibody levels are negative, the patient could have still had COVID19. While the assay is simple and can be read with the naked eye, there is more subjectivity to it compared to interpreting lines on lateral flow assays or light detection in chemiluminescent ELISA’s. Depending on mixing technique and provider comfort, there could be variability, likely necessitating training and/or deployment of controls in order to ensure proper reading of the assay. A negative control test without fusion protein will be important to include during clinical implementation for patients with positive tests, given that rare patients may have IgM autoantibodies causing agglutination.^31^ Given the similarities of this test to currently used ABO typing^32^ and Monospot assays for EBV antibodies,^33^ we ultimately do not see this being a significant barrier. Lastly, the fusion protein stability is unknown, which is important for use in low-resource countries. Past studies with similar fusion proteins for hemagglutination assays found stability at least 30 days at 37°C and 6 months at 4°C with no loss of assay agglutination activity.^14^ Ultimately, similar investigations will be important to conduct for each fusion protein employed to detect SARS-CoV-2.

For future directions, the design of the fusion protein can also be altered to enhance or detect certain subsets of SARS-CoV-2 proteins. For example, future iterations can employ a fusion protein of the entire or larger portions of the ectodomain of the spike protein, of which patients have been reported to have much higher antibody titers against.^28^ Moreover, peptides from the SARS-CoV-2 nucleocapsid protein can also be employed. Further testing should be able to reduce this incubation time by optimizing fusion protein and reagent concentrations. Ongoing studies will confirm the sensitivity and specificity of the assay with more COVID-19 patient samples. Similar prior assays for HIV demonstrated 100% sensitivity (n=94) and 99.5% specificity (n=596), suggesting similar efficacy for SARS-CoV-2 could be achieved.^34^

For clinical applications, the key utility of the test will be the ability to rapidly detect who has been exposed and has antibodies against COVID-19 in the population. Beyond epidemiology, any positive test for antibodies binding to RBD would be highly suggestive of the presence of neutralizing antibodies that would be protective of reinfection.^24^ The ability to detect potentially neutralizing antibodies in the plasma of COVID-19 recovered patients would be useful in the continued deployment of convalescent sera or hyperimmune globulin as a therapy against COVID-19.^35,36^ Convalescent serum has shown potential early promise against the SARS coronavirus,^37^ but studies are generally limited in efficacy by variable and low neutralizing antibody titers in donors.^38^ The ability to rapidly screen donors for high titer antibodies against SARS-CoV-2 could facilitate the scaling of convalescent plasma donation.^39^ Monitoring antibody development in patients may also help stratify COVID-19 patients and offer prognostic value for those who may clinically improve versus suffering respiratory demise.

In conclusion, our RBC agglutination assay with cross-linked viral antigen-antibody fusion may represent a simple, cheap and scalable way to dramatically increase our capability of detecting antibodies against SARS-CoV-2. It may find utility, particularly in low-resource settings, in the efforts to combat this pandemic. Additional work is needed to confirm and extend these findings.

## Acknowledgments

We would like to thank all the brave health care workers continuing to serve during the COVID-19 pandemic, as well as the environmental services staff at Johns Hopkins for continuing to come to work and clean labs, aiding in our pursuit of this goal.

This work was funded by the Johns Hopkins Department of Pathology Fred and Janet Sanfilippo Research Award (R.L.K), and partially supported by the National Institutes of Health, National Institute of Diabetes and Digestive and Kidney Disease grant R01DK106109 (Z.Z.W).

